# Chromatin compaction states, nuclear shape fluctuations and auxeticity: A biophysical interpretation of the epigenetic landscape of stem cells

**DOI:** 10.1101/419556

**Authors:** Kamal Tripathi, Gautam I. Menon

## Abstract

When embryonic stem cells differentiate, the mechanical properties of their nuclei evolve *en route* to their terminal state. Measurements of the deformability of cell nuclei in the transitional state that intervenes between the embryonic stem cell state and the differentiation primed state of mouse stem cells, indicate that such nuclei are *auxetic i.e.* have a negative Poisson’s ratio. We show, using a theoretical model, how this unusual mechanical behaviour results from the coupling between chromatin compaction states and nuclear shape. Our biophysical approach, which treats chromatin as an active polymer system whose mechanics is modulated by nucleosome binding and unbinding, reproduces experimental results while providing new predictions. We discuss ways of testing these predictions. Our model suggests a biophysical interpretation of the epigenetic landscape of stem cells.

## Highlights

- A mathematical model describes the coupling of chromatin compaction states with fluctuations in the shape of stem cell nuclei
- It explains the observation of auxetic behaviour in mouse stem cells in the transitional state
- The results agree with experiments and the model provides several testable predictions
- These results suggest a biophysical interpretation of Waddington’s epigenetic landscape in terms of chromatin compaction states

## Introduction

Embryonic stem cells (ES cells) occupy the apex of a hierarchy of cellular states (Young, 2011). They self-renew, maintaining their “stemness”, but can differentiate into varied cellular lineages when supplied appropriate biochemical or mechanical cues (Suda et al., 1987; Heo et al., 2018). Lineage choice results from shifts in patterns of gene expression, controlled by the rewiring of gene regulatory networks through the differential expression of transcription factors, but also from modifications to chromatin structure, such as through the methylation of cytosine residues in CpG dinucleotides by DNA methyl-transferases, the incorporation of histone variants, the post-translational modification of amino acid residues in histone sub-units and the action of structural proteins which alter chromatin conformation (Berger, 2007; Hawkins et al., 2010). Such epigenetic modifications modify the local structure and biophysical properties of chromatin.

Changes in patterns of gene expression should have biophysical correlates, since they require actively transcribed genes to be more accessible *vis a vis.* silenced genes (Rando and Chang, 2009). Regions of chromatin which see higher levels of transcriptional activity are typically more loosely packed than gene poor and relatively more compact heterochromatin regions, correlating to local epigenetic marks (Narlikar et al., 2002). In differentiated cells, trimethylation at the histone location H3K4 results in an open chromatin configuration characteristic of actively transcribed euchromatin, whereas condensed, inactive heterochromatin is enriched in H3K9 and H3K27 trimethylation (Du et al., 2015). ES cells are known to be transcriptionally hyperactive and to possess an open chromatin state with reduced levels of heterochromatin (Efroni et al., 2008). Such a state may contribute to the maintenance of pluripotency (Gaspar-Maia et al., 2011).

The distinction between locally more and less compact local chromatin configurations is clearly relevant to the biophysics of chromatin. It may assume added importance in the highly dynamic stem cell state (McNally, 2011). ES cell chromatin is known to be “hyperdynamic”, with histones binding and unbinding locally at an enhanced rate compared to differentiated cells (Meshorer et al., 2006). These increased fluctuations are accompanied by enhanced fluctuations in nuclear shape and size, arising largely from the absence of A-type lamins in the nuclear lamina of ES cells (Lammerding et al., 2004; Talwar et al., 2013). A biophysical link between nuclear mechanics, chromatin packaging and lineage choice is suggested by the observation that purely mechanical cues, such as substrate stiffness or substrate structure, are sufficient to drive stem cell differentiation into preferred lineages (Engler et al., 2006; Yim and Sheetz, 2012; Hwang et al., 2013).

Global chromatin remodelling occurs during differentiation, which results in a transition between an open chromatin configuration to a more compact state (Chen and Dent, 2014; Ugarte et al., 2015). Prior to lineage commitment, ES cells exhibit de-condensed chromatin and soft nuclei. A slowing down of histone dynamics and the stiffening of the nuclear envelope accompanies differentiation (Engler et al., 2006; Justin and Engler, 2011; Evans et al., 2009). The interplay of chromatin packaging with fluctuations of the relatively pliable chromatin-enclosing nuclear envelope might then reasonably be expected to underly the special biophysical properties of the stem cell state (Boskovic et al., 2014; Dado et al., 2012).

Waddington originally visualised the differentiation of stem cells in terms of a set of branching tracks representing different cell fate choices (Waddington, 1947). A subsequent, more pictorial version of this idea used the analogy of a ball rolling along an “epigenetic landscape” with minima chosen to represent stable differentiated states (Gilbert, 2000; Waddington, 2014). Stable positions in this landscape have been argued to correspond to attractors of a high-dimensional nonlinear dynamical system controlled by feedback (Huang, 2012). This provides a particularly appealing and pictorial way of understanding how stem cell differentiation into specific cell lineages can be visualized. Such ideas connect naturally to other landscape descriptions of bio-physical states and phenomena (Kauffman, 1992; Onuchic et al., 1997). However, the experimental corollaries of an epigenetic landscape and how, in particular, nuclear mechanics might enter its description, are little understood.

We ask whether biophysical measurements of the mechanical properties of stem cell nuclei can provide insights into these questions (Miroshnikova et al., 2017; Chalut et al., 2012; Swift and Discher, 2014). We first note that almost all materials have a positive Poisson’s ratio, becoming fatter in the transverse direction when compressed uniaxially along a longitudinal dimension (Landau and Lifshitz, 1986; Chaikin et al., 1995). Materials with a negative Poisson’s ratio, among them foams, are termed auxetics (Evans and Alderson, 2000; Grima et al., 2006). Pagliara *et al.* (Pagliara et al., 2014) report results from atomic force microscopy (AFM) measurements of the reduced modulus K = E/(1 *– v*^2^), with E the unaxial stiffness and *v* the Poisson’s ratio, of mouse embryonic stem cell (ESC) nuclei exiting the pluripotent state *en route* to differentiation. In this transitional state of embryonic stem cells (T-ESC), monitored through levels of the GFP-labeled pluripotency marker *Rex1* and obtained when specific inhibitors preventing the transition to a differentiation primed state are removed, the cell nuclei were noticed to become smaller by about 5-10% in cross section when compressed to the level of about 2 *μ*m with the AFM probe (Pagliara et al., 2014). Similar results were obtained by observing changes in nuclear dimensions when T-ESC cells were set in flow along a microchannel. Whereas both the naive embryonic stem cell state (N-ESC) as well as the differentiation primed state exhibit a positive Poisson’s ratio, the T-ESC that intervenes between them is thus auxetic, with a *negative* Poissons ratio. Pagliara *et al.* suggest that the auxetic phenotype might be connected to chromatin de-condensation, since chromatin in the transitional state is less condensed than in either the embryonic stem cell state or the differentiation primed state (Pagliara et al., 2014). Disrupting the actin cytoskeleton through Cytochalasin D treatment did not remove auxeticity, indicating that it might naturally originate in the biophysical properties of the nucleus itself and not of the extranuclear environment.

A biophysical approach to understanding this experiment and its larger implications, both for chromatin biophysics and the origins of stem cell behavior, must identify relevant variables of interest, especially those that are amenable to measurement. Fig. 1(a) shows a schematic of the experiments of Ref. (Pagliara et al., 2014) while Fig. 1(b) illustrates how the on-off dynamics of nucleosomes in the stem cell state might alter chromatin packaging. Fig. 1(c) illustrates the definitions of the fundamental mechanical variables that enter our model. We use a single variable, labelled, to describe nucleosome-induced compaction of chromatin. The variable can be thought of as representing the number of nucleosomes bound to chromatin at a given time, with the biophysical interpretation that a larger number of bound nucleosomes yields a more compact chromatin structure. The structural variables *R*_‖_ = *R*_0_ + δ*R*_‖_ and *R*_⊥_ = *R*_0_ + *δR*_⊥_ denote nuclear dimensions parallel and perpendicular to the direction of the applied force f, as shown. Fig. 1(d)-(e) illustrate how nuclei deform under force in both the non-auxetic (d) and the auxetic (e) case. Finally. Figs. 1(f) and (g) supply schematics of auxetic and non-auxetic response to a force f, in the variables, *R*_∥_, and *R*_⊥_. How to derive the schematics of Figs. 1(f) and (g), including the behaviour of in both the auxetic and non-auxetic cases predicted by the theoretical formulation, is the subject of this paper.

**Figure 1:**
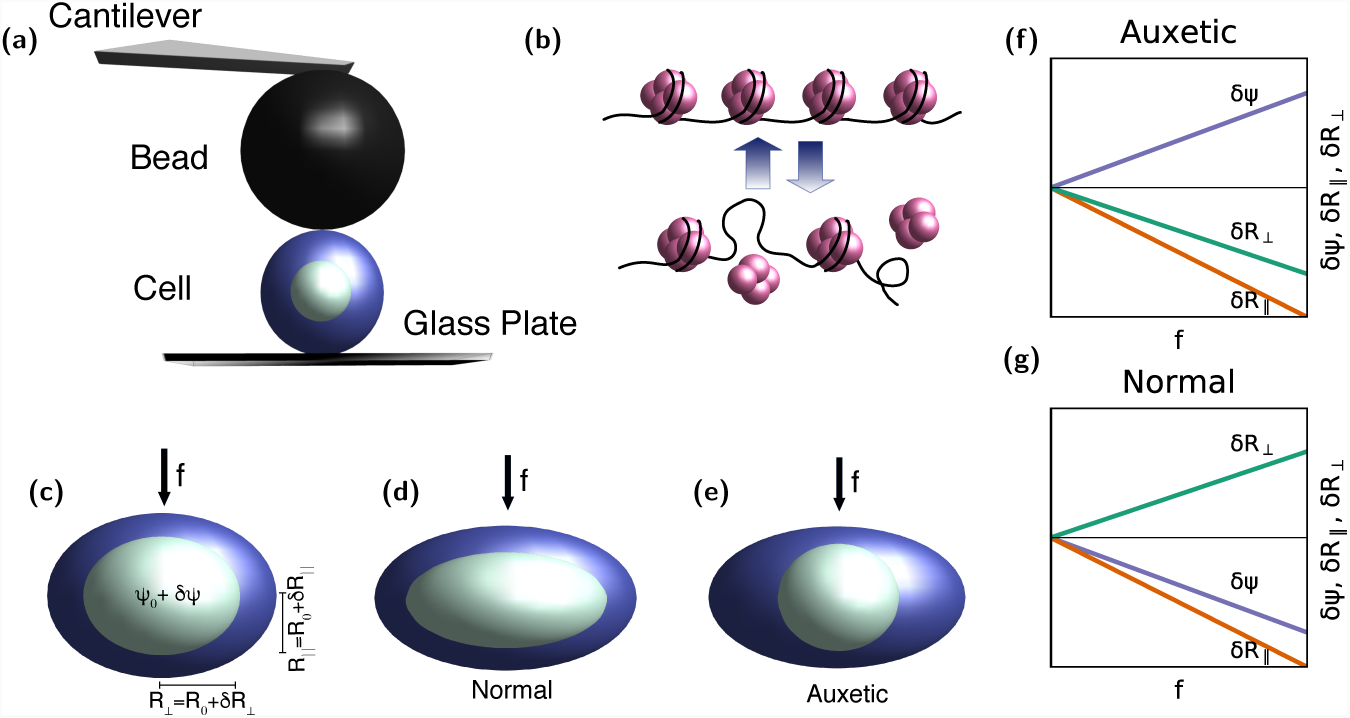
**(a)** Schematic of the AFM experiment of Ref. (Pagliara et al., 2014) **(b)** Fluctuations in chromatin compaction arising from the fast on-off dynamics of nucleosomes in the stem cell state, where histones are hyperdynamic **(c)** Definitions of the variables, *R*_∥_, and *R*_⊥_ in the AFM-based indentation experiment, including the applied force *f* arising from the indentation **(d)** Illustration of normal *i.e.* non-auxetic behaviour in the experiments, showing how the nucleus expands in the direction perpendicular to the applied force f, while the nuclear dimension in the direction parallel to the force contracts. **(e)** Illustration of auxetic behaviour, showing how the nucleus contracts both in the direction perpendicular to the applied force as well as in the direction parallel to it. The schematic plots in **(f)** for the non-auxetic case and **(g)** for the auxetic case show how the variables, *R*_∥_, and *R*_⊥_ behave in both limits as *f* is increased from zero. The unperturbed nucleus is taken to be spherical.

## Results

We propose that the coupling of nuclear size fluctuations with chromatin compaction states is responsible for auxetic behaviour. These variables takes the values Ψ_0_ and *R*_0_ *on average* but exhibit fluctuations around these values. Crucially, these fluctuations are not independent of each other. Large values of should represent large local compaction, as seen in the differentiated state, whereas small values indicate a more loosely bound, more permissive state. By considering only the overall compaction in Ψ, we are implicitly averaging over spatial variations in that quantity, a reasonable approximation when chromatin is more fluid-like than solid, as in ES cells.

Our model relies on four biophysical assumptions. These follow from the experimental observations. First, chromatin in the auxetic regime is compressible, a fundamental property of the auxetic state. Second, mechanical response to an external force in such a regime must be anisotropic. Such anisotropy is essential to auxetics, although it should not be intrinsic to isolated stem cells in the absence of an applied force. Third, a number of experiments indicating chromatin fluidity in all but terminally differentiated states argue that chromatin is best described as a confined, active polymer fluid in a semi-dilute regime (Pajerowski et al., 2007). (Indeed, the formation of heterochromatin foci has been discussed in analogy with active phase separation in liquid-liquid mixtures (Larson et al., 2017; Strom et al., 2017).) An alternative view of auxeticityy which considers the nucleoplasm to be a gel and uses ideas from phase separation is described in Ref. (Yamamoto and Schiessel, 2017). We will treat activity as equivalent to a (higher) effective temperature (Ganai et al., 2014; Agrawal et al., 2017).

Fourth and finally, we assume that auxetic behaviour arises from the form of the coupling of chromatin compaction states to mechanical variables, which we choose as nuclear dimensions parallel to, as well as perpendicular to, the applied force. These four assumptions, all reasonable from a bio-physical standpoint, inform our mathematical model. We use them to derive a model non-linear dynamical system describing auxetic behaviour in the transitional state of stem cells.

## Model description

Our equations are formulated in terms of δΨ, *δR*_∥_ and *δR*_⊥_ defined as in Fig. 1 and discussed in Methods. The equations describing how these quantities change in time take the form

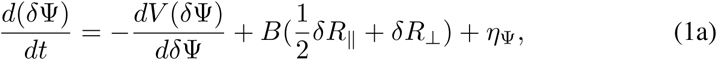

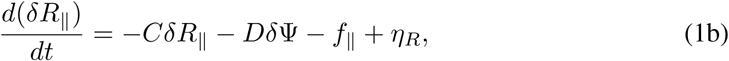

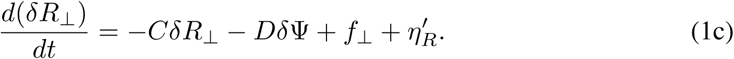

We have used our freedom to choose units suitably to “de-dimensionalize” the coefficients that appear in these equations. The first term on the right-hand side of each of these equations represents the independent relaxation of fluctuations away from {*Ψ*_0_, *R*_0_}. We assume that the δΨ variable relaxes subject to an effective “epigenetic” potential V (δΨ). The interpretation of this term as describing an epigenetic potential will become clearer as we proceed. The second term couples δΨ to the mechanical variables *R*_∥_ = *R*_0_ + *δR*_∥_ and *R*_⊥_ = *R*_0_ + *δR*_⊥_, with coefficient B; the relative factor of 2 accounts for the 3-d geometry. This is the simplest linear form that these equations can take. Their biophysical content lies in the estimates of the numerical values associated with the coefficients. More subtly, the coupling between chromatin compaction and nuclear dimensions is to be found in the cross-terms in Eq. 1.

In the absence of a force, *R*_∥_ and *R*_⊥_ are equivalent. The symmetry between them is broken only by f_*∥*_ and f_*⊥*_. These forces represent both external forces as well as forces that arise from the remodelling of the extranuclear actin cytoskeleton, which can be assumed to be uniform in if f_*∥*_ = 0. We can assume that f_*∥*_ couples primarily to *δR*_∥_ whereas f_*⊥*_ couples to *δR*_⊥_. In the absence of external forces, the two equations reduce to a single one. The quantity *C* represents a ratio of time-scales for the relaxation of the and the *R* variables. If Ψ_0_ represents a stable state, or at least a state that evolves slowly on the time-scale of the fluctuations δΨ, we can expand in these fluctuations. At the simplest level then, these fluctuations are subject to a harmonic potential. The case where *δR*_∥_ = *δR*_⊥_ *≡ δR*, with f_*⊥*_ = f_*∥*_ = 0 and the V (δΨ) term chosen to be bistable, was studied in Ref. (Talwar et al., 2013), in the context of nuclear size oscillations in the ES state of mouse stem cells. We will use this more specific form of these equations when we identify signals of auxetic behaviour in fluctuations within the undeformed steady state. Our results suggest that signatures of the transition between auxetic and non-auxetic behaviour might be most easily seen in these fluctuations.

In Ref. (Talwar et al., 2013), in a description of enhanced fluctuations in mouse N-ESCs, B > 0 was assumed. The physical interpretation there was that *increasing* the size of the nucleus would expose binding sites for histones. This leads to a concomitant *increase* in *ϕ* which would then drive the nucleus to shrink (Talwar et al., 2013). The coupled dynamics of the fast histone on-off rates in the hyperdynamic case with the slower fluctuations in nuclear size leads to interesting fluctuation behaviour. Such a choice of sign leads inevitably to non-auxetic behavior; see below.

Experiments show that chromatin is most decondensed in the transitional state, as opposed to either the ES state or the differentiation primed states between which it intervenes (Pagliara et al., 2014). A further expansion of nuclear dimensions might then be expected to result in the expulsion of nucleosomes, rather than their accumulation, in this intermediate state. Incorporating this into the modelling requires that we consider the case where *B* < 0. Indeed, treating the naive pluripotent ESC state with trichostatin A, an HDAC inhibitor that globally decondenses chromatin, made the N-ESC auxetic, arguing for the connection our modelling proposes. We can thus view the transition between the naive stem cell, the transitional state and the differentiation primed state in terms of a re-entrant behaviour in the sign of *B*. This is an experimentally testable prediction.

The on-off dynamics of histones is inherently noisy. Our equations account for such stochastic effects, represented as additive noise with standard properties, with terms represented by *η*_Ψ_, *η_R_* and 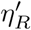. In general, the effects of the noise should be most significant for the fast fluctuating variable. We thus choose to retain only the Gaussian-distributed, delta-correlated *η*_Ψ_ term in our equations, setting 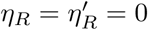.

### Auxetic and normal mechanical behaviour in a model description of nuclear indentation

The AFM indentation experiment corresponds to taking *f* = f_*∥*_ ≠ 0, setting f_*⊥*_ = 0. The set of model equations, Eqs. 1 have a number of parameters, which we fix using experimental and theoretical input. The choice of parameters and the range of values they can take are discussed in Methods. The solutions of these equations are provided in Supplementary Information.

Fig. 2 shows plots of *δ*Ψ, *δR*_∥_ and *δR*_⊥_ (Fig 2, panels (e) -(h)) for small *f*, as obtained from our model equations. The quantities *δ*Ψ, *δR*_∥_ and *δR*_⊥_ vary linearly with *f*, a consequence of the fact that we assume that *V* (δΨ) increases quadratically about its stable value. This is across the parameter values shown in Fig. 2, panels (a) - (d), for the state points (B,C) marked on the figures with the filled black circle. These plots are for a choice of parameters corresponding to non-auxetic i.e. regular behaviour. For the normal *i.e.,* non-auxetic state, the slope of *δR*_∥_ and *δR*_⊥_ *vs. f* should have opposite signs.

Our model predicts that the slope of *δ*Ψ *vs. f* is negative *i.e.* the compaction decreases with the applied force in the non-auxetic state; see Fig. 2, panels (e) and (h).

In Fig. 2, panels (m) - (p), we also show results for the auxetic case, where the slope of *δR*_∥_ and *δR*_⊥_ *vs f* have the same sign. Note that *δR*_∥_ and *δR*_⊥_ now both decrease with f. This indicates auxetic behaviour. This behaviour is seen across the parameter values shown in Fig. 2, panels (i) - (l), for the state points marked on the figures with the filled black circle. These parameter values are chosen in the regime where the fixed point is stable, shown in blue. (The grey region shows the regime in which the equations have unstable solutions.) In the auxetic case, the slope of *δ*Ψ *vs. f* is positive.

**Figure 2:**
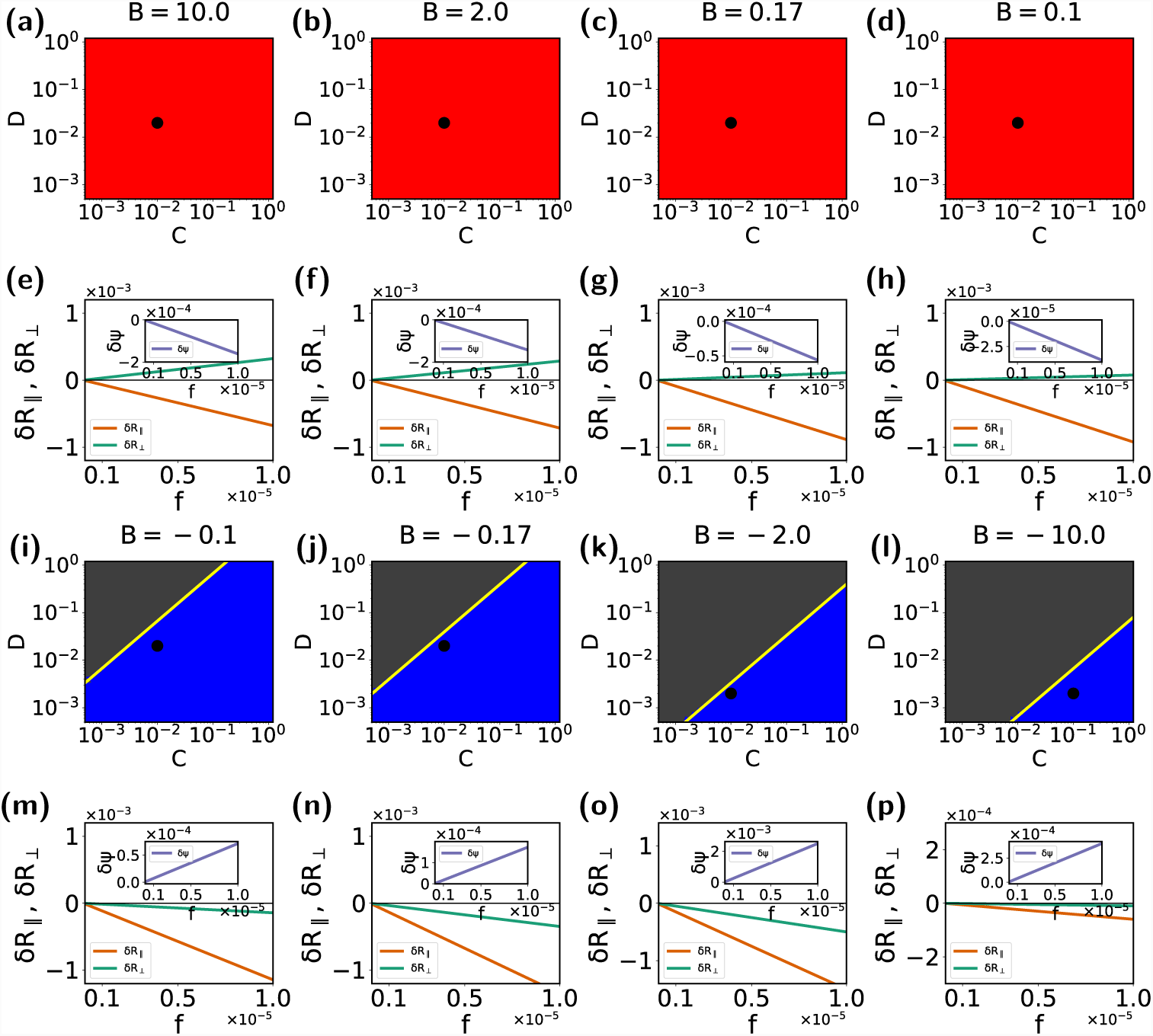
Parameter choices for C and D and: **(a)** B = 10.0, **(b)** *B* = 2.0 **(c)** *B* = 0.17 **(d)** *B* = 0.1, all in the regime of non-auxetic (regular) behavior. The behaviour of the dynamical variable with the increase in force for the parameter values **(e)** *C* = 0.01, *D* = 0.02, *B* = 10.0. **(f)** *C* = 0.01, *D* = 0.02, *B* = 2.0. **(g)** *C* = 0.01, *D* = 0.02, *B* = 0.17. **(h)** *C* = 0.01, *D* = 0.02, *B* = 0.1. Parameter choices for C and D, in the auxetic regime with **(i)** *B* = – 0.1, **(j)** *B* = – 0.17 **(k)** *B* = – 2.0 **(l)** *B* = 10.0. The line separating blue and gray regions marks the stable-unstable boundary. The behaviour of the dynamical variable with the increase in force for the parameter values **(m)** *C* = 0.01, *D* = 0.02, *B* = –0.1. **(n)** *C* = 0.01, *D* = 0.02, *B* = –0.17. **(o)** *C* = 0.01, *D* = 0.02, *B* = –2.0. **(p)** *C* = 0.1, *D* = 0.02, *B* = –10.0. (Red and blue colours in the colour plots show the regions where a stable solution is obtained (red = normal, blue = auxetic) while the grey colour shows where solutions become unstable).

Thus, the solutions of our model equations yield both auxetic and non-auxetic behaviour, controlled by the sign of *B* in Eqs. 1. The results are consistent with the schematics of Fig. 1 (f) and (g), which show how chromatin compaction varies upon the application of an external force. The additional information they provide relates to the behaviour of the compaction variable. As shown in Methods, the parameter values we derive are consistent with experimental measures of auxeticity in transitional stem cell nuclei.

### Describing nuclear shape changes in micro-channel flow

Nuclear indentation through the AFM method described in Ref. (Pagliara et al., 2014) provides a direct way of accessing the auxetic mechanical behaviour of the nucleus. Here, a fixed force is applied along the longitudinal (*k*) direction and a transverse (⊥) deformation measured. An alternative method involves an optofluidic assay, in which cells are passed through narrow micro-channels of controlled width. These cells are then imaged through fluorescence microscopy of Syto13 labeled cells. When the width of the channel is comparable to the cell size, this constrains cell dimensions. A further complication is the role of stretching stresses caused by cytoskeletal strain acting when cells are confined to the micro-channel. Given our model assumptions, we may model the confined case by accounting both for *f*_*∥*_ and *f*_*⊥*_ in the governing equations, Eqs. 1. Whereas *f*_*∥*_ is primarily controlled by the size of the constriction through which these cells pass, *f*_*⊥*_ derives from the anisotropic remodelling of the actin cytoskeleton.

The geometry of the micro-channel experiments is shown in Fig. 3(a), where we show a cell confined to a channel whose width is comparable to cell dimensions. In the experiments the channel width is 12 *μ*m while cell sizes range from 6 *μ*m to 14 *μ*m. Fig. 3(b) shows a schematic of the effects of the combination of longitudinal and transverse forces applied to cells of different sizes, as obtained in our calculations; see below. For small cells in the auxetic case, if they have unconfined dimensions much smaller than the channel width, their longitudinal and transverse dimensions *increase* when they are confined to the channel. For larger cells in the same limit, both dimensions *decrease*. These are consistent with expectations from auxetic behaviour. On the other hand, irrespective of cell sizes in the non-auxetic case, the longitudinal dimension *decreases* while the transverse dimension *increases*. These are consistent with the behaviour shown in Fig. 3(b). These schematic results recapitulate the results of Ref. (Pagliara et al., 2014).

**Figure 3:**
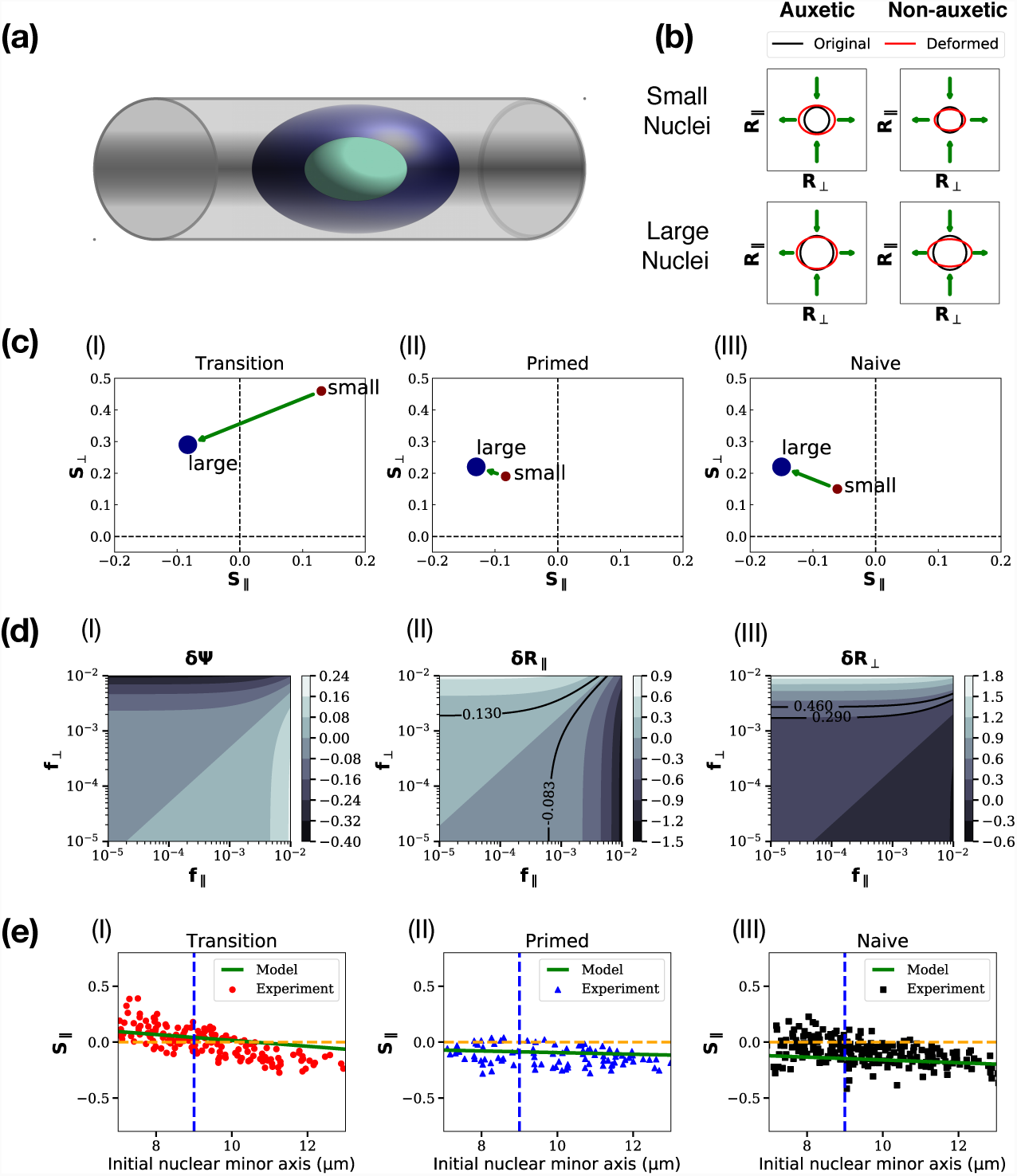
**(a)**: Schematic of a cell confined to a microchannel with width comparable to cell dimensions, **(b)** schematic of the effects of the combination of longitudinal and transverse forces applied to cells of different sizes. These follow from our calculations and are consistent with the results of Ref. (Pagliara et al., 2014), **(c)** Plots of *S*_∥_ and *S*_⊥_ extracted from experiments, for the transitional, primed and naive ES cell states. The arrow connects the two terminal points. **(d)** Contour plots for δΨ, *δR*_⊥_ and *δR*_∥_, against (*f*_∥_, *f*_*⊥*_), with solid lines showing loci of constant strain **(e)** Predictions for transitional, primed and naive ES cell states, of *S*_∥_ and *S*_⊥_. The straight line represents experimental predictions for intermediate cell sizes. Experimental data are digitized from the scatter plot of Fig. S10 of Ref. (Pagliara et al., 2014) and shown on the same figure

To extend this to the mechanical response of stem cells of various sizes in a microchannel, our modelling strategy is the following. The experiments, performed on a range of cell sizes at a fixed micro-channel width, obtain longitudinal and transverse strains for an ensemble of cells of different sizes. At the extreme limits of cell sizes, Fig. S10 of Ref. (Pagliara et al., 2014) shows averaged strains in the parallel and perpendicular directions. These are proportional to *R*_∥_ and *R*_⊥_ in our definitions in Eq 1, and using *R*_0_ as our unit of length converts this proportionality to an equality. We label these strains as *S*_∥_ and *S*_⊥_ and display them in Fig. 3(c) (I) - (III), for the transitional, primed and naive ES cell states. Starting with these results, we can invert the relationship between strains and forces, finding the effective *f*_*∥*_ and *f*_*⊥*_ that produce these strains.

We can now explore the space of values of (f_∥_, *f*_*⊥*_), constructing contour plots of δΨ, *δR*_⊥_ and *δR*_∥_, as shown in Fig. 3 (d) (I) - (III). The parameters chosen are for the smallest and the largest cells, using the data shown in Fig. 3(c) (I). The solid lines in Fig. 3 (d) (II) - (III) represent a choice of a few lines of constant strain in each case, as a function of *f*_*∥*_ and *f*_*⊥*_. These lines then *predict* the forces (f_∥_, *f*_*⊥*_) required to create a fixed strain across cells of different sizes.

The extremal points of Fig. 3(b) (I) - (III) are now associated to points on the (f_∥_, *f*_*⊥*_) surface. We can then model the data for cells of sizes intermediate from these by supposing that *f*_*∥*_ and *f*_*⊥*_ vary independently and linearly between their terminal values. We ask if these results can fit data for intermediate cell sizes, shown in the scatterplot illustrated in Fig. 3 of Ref. (Pagliara et al., 2014). These results are shown in Fig. 3(e) (I) - (III), for transitional, primed and naive ES cell states. The experimental data are shown as points while the theoretical prediction that follows from our analysis is shown as the green line. In all three cases, there is an approximate linear relationship between *S*_∥_ and S_⊥_ that our calculation captures. The magnitude of the strains at intermediate values of cell size is correctly rendered.

Our model thus, despite its simplicity, captures all essential features of the data of Ref. (Pagliara et al., 2014). As we have pointed out, the model can then be used to provide specific predictions for mechanical response in cells of different sizes. Also, even though the chromatin compaction variable δΨ was not measured in those experiments, our model provides specific predictions for how this quantity varies across different cell sizes in comparison to the width of the microchannel. This prediction is experimentally testable.

### Autocorrelations and cross-correlations of chromatin compaction and nuclear dimensions in the auxetic regime

The previous sections explored the use of an external force, either applied directly using an AFM tip or indirectly by confining cells to a narrow microchannel, in understanding auxetic and non-auxetic behaviour. However, our general model formulation suggests how less invasive ways of probing the coupled mechanical response of chromatin compaction and nuclear dimensions might provide useful information. Let us assume that we can measure both chromatin compaction as well as the dimensions of the nucleus simultaneously as a function of time - possible ways of doing this are discussed later. Assuming an initially spherical nucleus, *R*_∥_ and *R*_⊥_ coincide, since now there is no externally imposed direction that leads to an anisotropic mechanical response. The only relevant mechanical variable is then *R*(*t*), the radius of the spherical nucleus as a function of time. Our equations are now simpler, since they involve only the two variables, Ψ and *R*. The solution to the full set of equations is provided in Supplementary Information.

**Figure 4:**
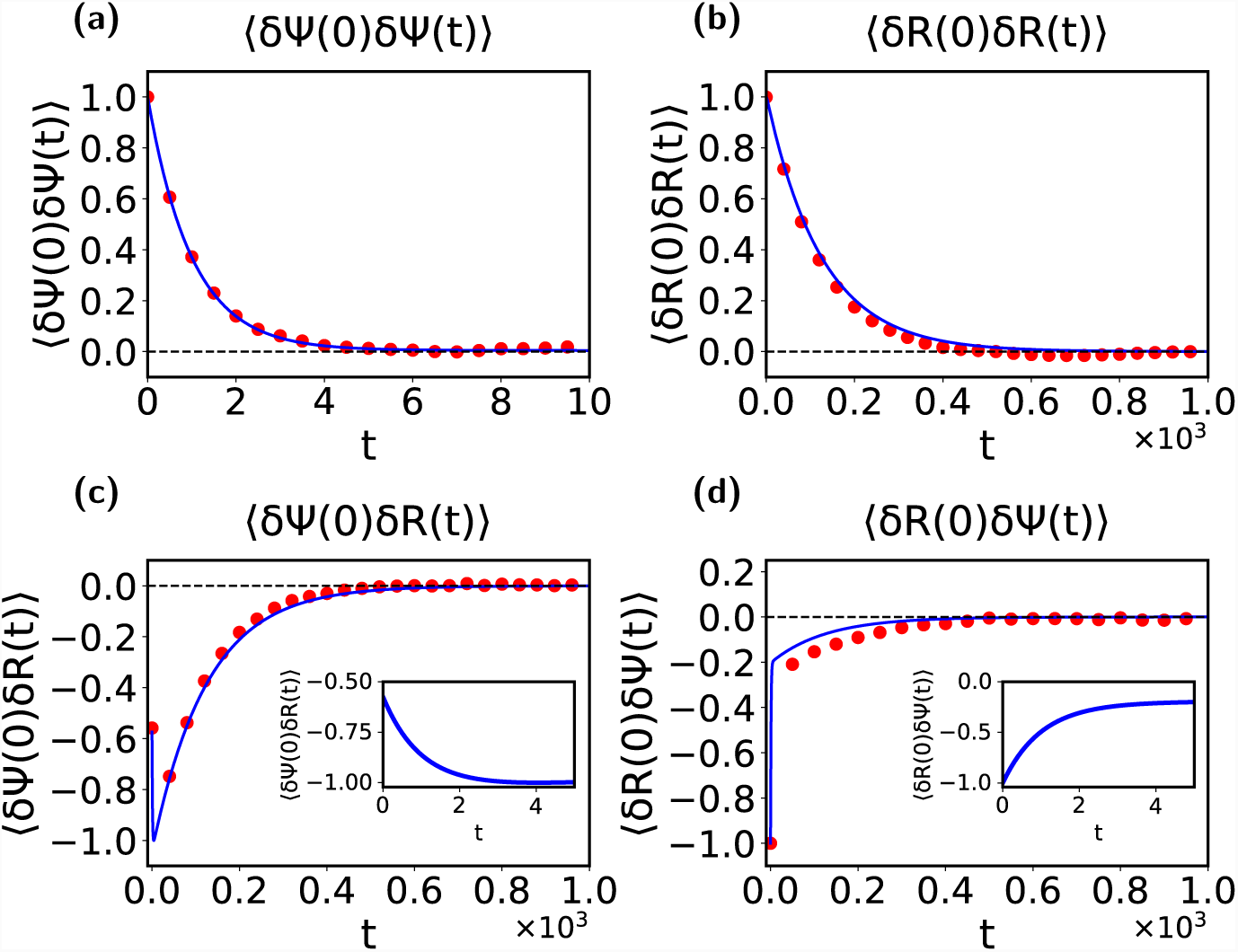
Computations of autocorrelations and cross-correlations in the simplified 2-component model, in the auxetic regime, with *C* = 0.01, *D* = 0.02 **and *B*** = –0.17. We illustrate the calculation of the following correlation functions: **(a)** The autocorrelation of the *δ*Ψ variable 〈 *δ*Ψ (0)*δ*Ψ (t)〉, **(b)** The autocorrelation of the *δR* variable, 〈*δR*(0)*δR*(t)〉, **(c)** The cross-correlation between δΨ and 〈*δR*, δΨ (0)*δR*(t), **(d)** The cross-correlation of *δR* and δΨ, *δR*(0)δΨ (t)〉. The insets show the behaviour close to the origin in two special cases where there is a competition between the two time-scales for relaxation. Points represent the numerical solution of the Langevin equations while lines represent the analytic formulae.

Given measurements of (*t*) and *R*(*t*) = *R*_0_ + *δR*(*t*), we can ask whether signatures of auxetic and non-auxetic behaviour might be visible in such measurements. Since such measurements provide data in time, we can compute autocorrelations of these variables as well as their cross-correlations. The solutions to these equations can be computed explicitly and are provided in Supplementary Material. Fig. 4 shows our computation (lines: exact calculations; points: numerical solutions of the Langevin equation) of autocorrelations and cross-correlations in the model, with parameters chosen within the auxetic regime.

The autocorrelations 〈*δ*Ψ (0)*δ*Ψ (t)*〉* and *〈δR*(0)*δR*(*t*)*〉* are shown in Fig. 4(a) - (b) whereas the cross-correlations 〈δΨ (0)*δR*(t)〉 and 〈δΨ (0)*δR*(t)〉 are shown in Fig. 4(c) - (d) respectively. The insets expand the behaviour of the cross-correlation functions close to the origin, where two time-scales for relaxation compete. The time-scale for the relaxation of autocorrelations in the variable is substantially smaller than for the *R* variable. The cross correlations 〈δΨ (0)*δR*(*t*)〉 and 〈δΨ (0)*δR*(*t*)*〉* both relax to zero in an interesting two-step way, with a sharp initial step reflecting the relaxation of the fast variable followed by a slower relaxation, primarily driven by the *R* variable.

We can further investigate (see Supplementary Information) model predictions for the case in which a weak force is applied and allowed to vary in time in a sinusoidal fashion. For our linear system of equations, this then implies that quantities such as Φ, *R*_∥_ and *R*_⊥_ should also oscillate at the same frequency, but with a phase lag between them. This phase lag predicts the relative importance of what is termed reactive and dissipative response, with the first largely located in the elastic properties of the nuclear envelope and the second associated to dissipation connected to the flow of fluid across the nuclear envelope as well as of the friction encountered by chromatin as its fluctuations relax. These can be predicted from the theoretical formulation, and indeed are the focus of standard experiments in the physics literature that studies soft materials, but whether their experimental analogue can be probed in biophysical measurements on stem cells is an open question.

Extracting behaviour as shown in Fig. 4 would constitute a powerful test of model predictions.

### Correlations across the auxetic-nonauxetic boundary as probes of the transition

Our model describes chromatin compaction states using a single variable Ψ, with larger values of Ψ representing overall more compact states of chromatin packing. We suggest that Ψ fluctuates in time about an approximately constant value, but that these fluctuations are constrained by an “epigenetic potential”, defined as V (δΨ), that controls how large they can be. These fluctuations are also constrained by their coupling to nuclear dimensions through the variables *δR*_∥_ and *δR*_⊥_. They are influenced, as well, by the inherent noisiness of nucleosome on-off dynamics in a hyperdynamic state. All these effects are included in our model.

This choice of an “epigenetic potential” identifies the relevant *biophysical* distinction between more open, gene-rich euchromatin and more tightly bound, gene-poor heterochromatin as broadly being one of local compaction. We project the multidimensional landscape of potential chromatin states that Waddington envisaged, which should be more generally describable through a spatially varying and sequence-dependant compaction field, onto a single scalar compaction variable. Our equations then provide a way of understanding how such a compaction variable couples to mechanical variables describing nuclear shape and size.

How should we think of the epigenetic landscape in operational terms? Our results suggest a simple method for determining the location of the auxetic-to-non-auxetic transition. We will work in the limit described in the previous section, where we infer the transition by monitoring the system non-invasively, measuring only the variables Ψ and *R* as functions of time in steady state. From these measurements, we can calculate their autocorrelations and cross-correlations.

In Fig. 5, we show plots of the correlation functions *〈δR*(0)*δ*Ψ (t)*〉* and *〈δ*Ψ (0)*δR*(*t*)*〉*. These illustrate that *〈δR*(0)*δ*Ψ (*t*)*〉* is a good indicator of the transition from auxetic to non-auxetic behaviour, with *〈δR*(0)*δ*Ψ (*t*)*〉* changing the sign of its slope upon approaching its asymptotic value across the auxetic to non-auxetic boundary. On the boundary, there is no correlation at all, to this order, between fluctuations in the nuclear dimension and fluctuations in chromatin compaction. Since the change from auxetic to non-auxetic behaviour is marked by the parameter *B* changing sign, it must cross zero at least at one point. (Since the experimental sequence encountered as ES cells differentiate is: non-auxetic *→* auxetic *→* non-auxetic, this suggests that *B* should change sign at least twice. This is a specific prediction that can be addressed in experiments, as we discuss below.)

Now note that at this special point, fluctuations in decouple from fluctuations in the nuclear size variable to linear order; fluctuations in influence fluctuations in *δR* but not *vice versa*. As we show below, this provides a practical way of accessing the epigenetic potential V (*δ*Ψ).

**Figure 5:**
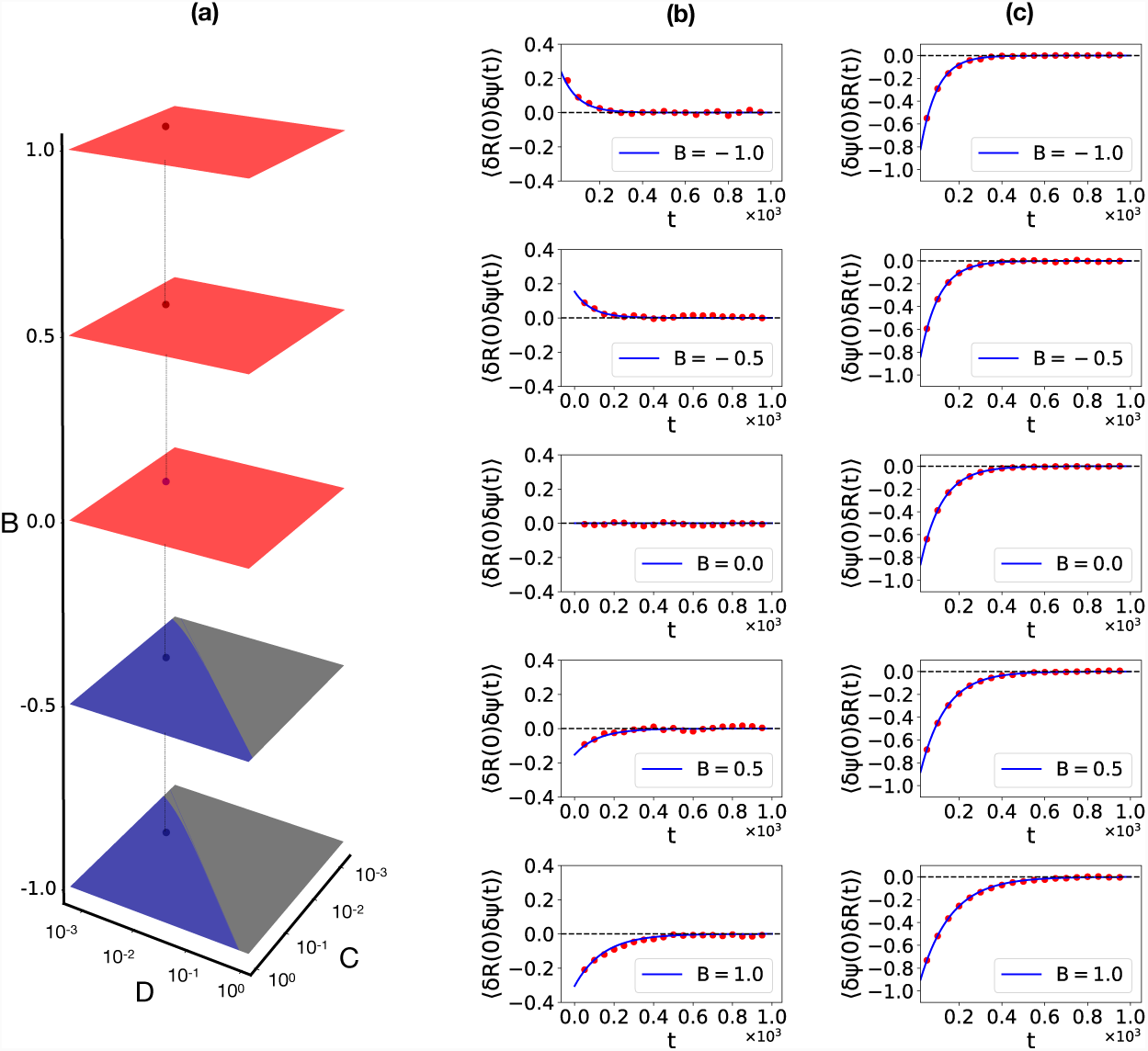
The left columns **(a)** shows our choice of parameters in (B,C,D) space, with B, shown on the vertical axis, varied so as to cross the auxetic to non-auxetic boundary. The two columns on the right, columns **(b)** and **(c)** illustrate the correlation function *δR*(0)*δ*Ψ (*t*) and *δ*Ψ (0)*δR*(*t*). Across the auxetic to non-auxetic boundary, where the sign of *B* changes, the variable decouples, at lowest order, from the *δR* variable, leading to a flat behaviour of the correlation *δR*(0)*δ*Ψ (*t*). In contrast, while is not influenced by *δR*, fluctuations in do couple to *δR*, leading to a non-trivial relaxation of the correlation function δΨ (0)*δR*(*t*). This change of sign of d *δR*(0)*δ*Ψ (*t*) /*d*t indicates that the auxetic to non-auxetic boundary has been crossed.

### Inferring an epigenetic potential from experimental data

We can describe the transition between ESC, T-ESC and differentiation-primed states in terms of a trajectory in the space of the variables *C, D* and *B*. As is standard, we can assume that the parameters controlling these variables must vary smoothly, since they reflect continuous shifts in the transcriptome; indeed the assumption of smooth variation is central to landscape ideas. As the stem cell transits between these states, it encounters auxetic (*B* < 0) [Fig. 6 (g) - (i)] and non-auxetic (*B* > 0) [Fig. 6 (d) - (f)] states, with an intervening *B* =0 state, [Fig. 6 (a) - (c)].

For each of these chosen values of *B*, we illustrate the choice of a specific epigenetic potential that we can model as a smooth function, shown via the solid lines in each sub-plot. We choose these functions to be (1) a simple quadratic potential, (2) a quartic potential with a shallow minimum at the origin and two symmetrically placed deeper minima on either side, as well as (3), the more complex case of a quadratic potential with a superposed sinusoidal modulation that provides more intricate structure. We do not yet know what form such a potential takes in the experiments, but intend to illustrate a method by which information from the measurement of fluctuations could help in its extraction.

For *B* = 0, as shown in Fig. 6 (a) - (c), given that *δ*Ψ (*t*) reflects its relaxation in the epigenetic potential, we form a histogram of *δ*Ψ values. Since the governing equation for the *δ*Ψ variable can be interpreted as a Langevin equation for a particle moving in the specified potential, the steady-state probability distribution of *δ*Ψ can be inferred from this histogram in a straightforward manner, as discussed in Supplementary Material. Figs. 6 (a) - (c) shows results from a numerical and analytic reconstruction of the effective assumed potential *V* (*δ*Ψ) using such a method. In this way, we thus proceed from the histogram of measured values to the epigenetic potential that controls such fluctuations. While the data used in these figures is “synthetic”, the procedure for extracting the potential from them is robust.

**Figure 6:**
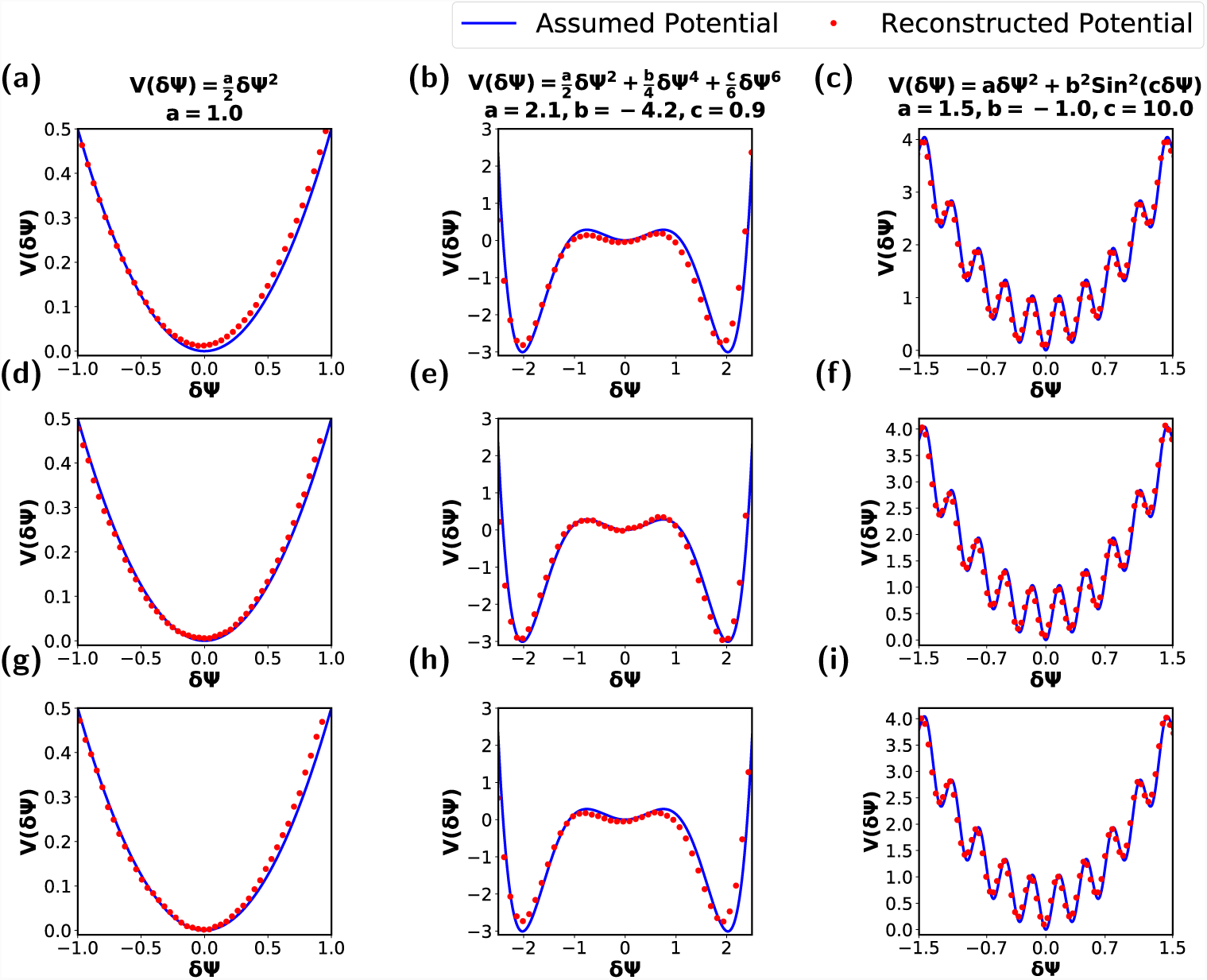
Reconstruction of potential landscape *V* (*δ*Ψ) **in the simplified 2-component model**, assuming a quadratic potential for **(a)** *B* = 0.0 **(b)** *B* = 1.0 *B* = -1.0, assuming a sextic potential for **(d)** *B* = 0.0 **(e)** *B* = 1.0 **(f)** *B* = -1.0 and assuming a quadratic potential with a superimposed sinusoid: *V* (*δ*Ψ) = *aδ*Ψ ^2^ + *b*^2^*Sin*^2^(*cδ*Ψ), for **(g)** *B* = 0.0 **(h)** *B* = 1.0 **(i)** *B* = -1.0

Provided *|*B*|* ≠ 0 is not too large and for parameter values comparable to the ones we use, this procedure reconstructs *V* (*δ*Ψ) reasonably well, a consequence of the fact that fluctuations in *δR* couple relatively weakly to fluctuations in *δ*Ψ. Figs. 6 (d) - (e), shows *V* (*δ*Ψ), obtained from histograms of *δ*Ψ values as in Figs. 6 (a) - (c), but for *B* = -1 and (e) *B* = 1. These suggest that one need not precisely locate the region where *B* vanishes for this approach to be of use.

## Discussion

An “epigenetic landscape”, whose lowest points represent gene expression patterns encoding specific differentiated states, is often pictorially represented in the following way (Huang, 2012; Furusawa and Kaneko, 2012): Imagine projecting all possible gene expression states onto a two-dimensional (XY) plane. This projection is constrained by the requirement that two nearby state points represent closely related expression patterns. (Naively, the rewiring of gene-regulatory networks required to convert expression programs from one cell type to another should be smaller the more similar these cell types are (Huang, 2012).) The height of a surface (the landscape) above a point on this plane can then be assigned to the relative “energy” of the state described by that point. The shape of the surface can then be used as a qualitative way of describing barriers to accessing different gene expression patterns starting from a given initial state.

The plasticity required of gene-regulatory networks in the stem cell states implies, in this pictorial analogy, that the shape of the landscape should determine which states will become unstable - and to which other states - as biochemical and mechanical parameters are changed. Biochemical parameters here could refer to levels of protein factors that modulate stemness while mechanical parameters could represent the stiffness and anisotropy of the substrate on which these cells are cultured (Li et al., 2012). If one imagines, as Waddington did, a ball rolling on this landscape as representing the stem cell state choosing between terminally differentiated states, the motion of the ball should be biased by the underlying shape of the landscape, including its peaks, ridges and valleys. The resulting energy surface can be depicted as a geographical landscape, along the lines of Waddington’s original picture. Such a qualitative picture also suggests that this landscape might also be thought of as *dynamic*, tilting and deforming to favour one set of states over others. This would then describe how an initial state might be guided to a specific cell fate as the cell integrates external environmental signals when driven to differentiate.

**Figure 7:**
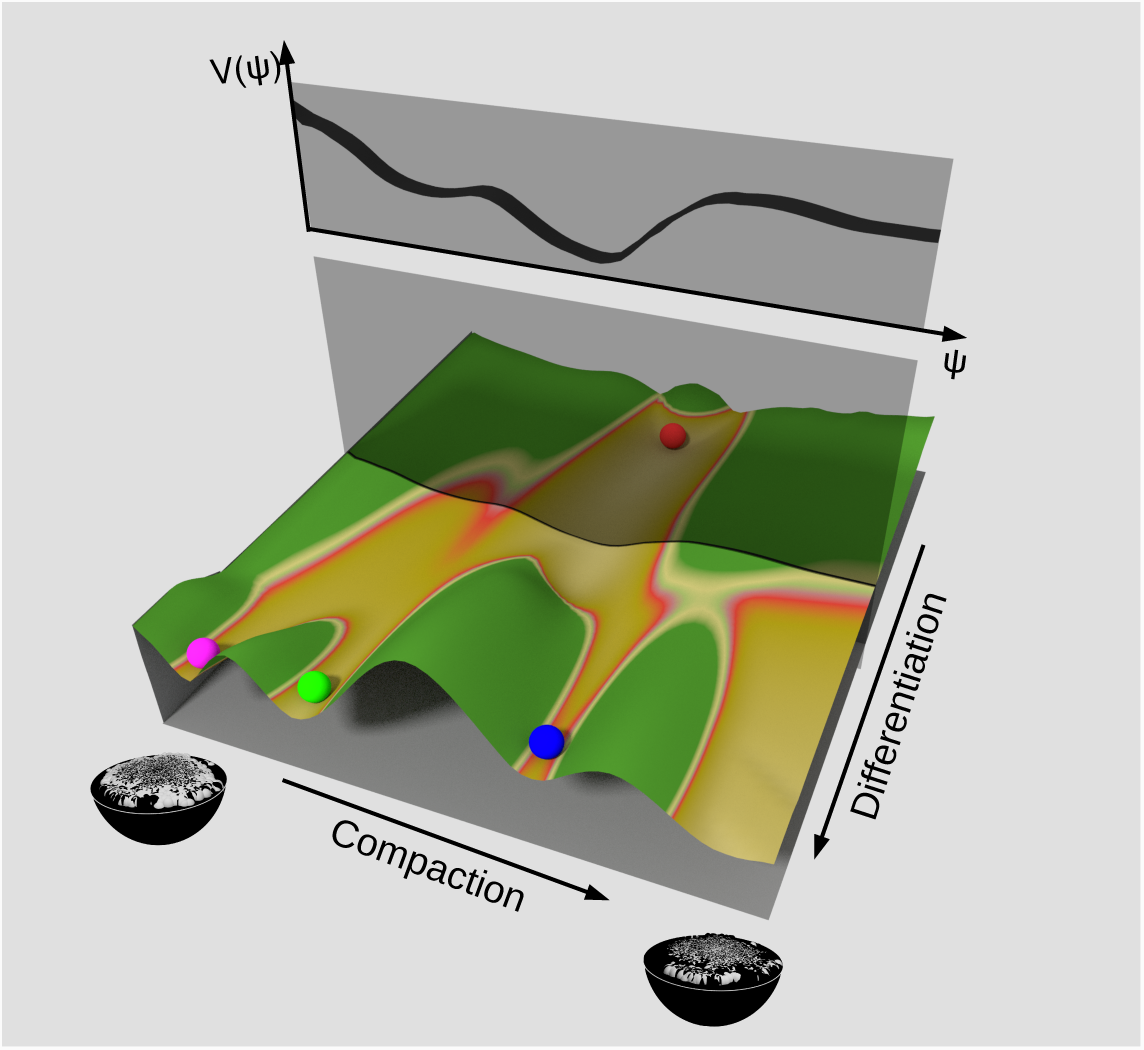
Schematic of Epigenetic Landscape in a Compaction variable. A pictorial representation of the epigenetic landscape, projected onto a single variable describing overall compaction. Points towards the back of the figure represented the ES cell state while points in the valleys towards the foreground represent differentiated states. As one moves from back to front, the figure describes how the effective potential governing overall compaction can be described via a cut through the landscape as shown.

The description of the previous paragraph proceeded along conventional lines. Our view here emphasizes biophysical aspects of this argument. Instead of projecting states depending on their proximity in gene expression space, we imagine them to be projected according to their level of chromatin compaction; arguments concerning the proximity of closely related cell types in such a “chromatin compaction” space should parallel those in the case of the “gene expression” space. To motivate this, we note that the relative ratio of heterochromatin to euchromatin varies across differentiated cell types (Rivera and Ren, 2013). It has been suggested that chromatin density might itself act to regulate gene expression in a stem cell population (Golkaram et al., 2017). While the embryonic stem call state has a chromatin organization that is best described as a highly correlated fluid, the differentiated state fluctuates far less, with condensed heterochromatin foci forming during the differentiation of pluripotent embryonic stem cells (Mao et al., 2015). The formation of heterochromatin domains has recently been argued to be mediated by phase separation (Larson et al., 2017; Strom et al., 2017). Together with the accumulation of silencing histone marks, this results in differential expression (Meshorer and Misteli, 2006). Classifying the epigenetic states underlying these cell types through their levels of local chromatin compaction should provide one approximate way of connecting the theoretical ideas presented here to experimental data.

This idea is illustrated in Fig. 7, which shows a schematic of such a landscape. The coloured balls towards the front of the figure represents stable, differentiated states. The ball at the back represents the ES cell state. As the cell differentiates, one imagines that the landscape is titled forward so as to allow the ball to fall towards these stable states. All possible accessible intermediate states can be represented, again pictorially, in terms of a plan that intersects this landscape. The curve that defines where these two curves intersect provides a one-dimensional surface, to be identified with the V (δΨ) of our discussion.

We stress that *all* projections from a high-dimensional space to a low-dimensional one, involve a loss of information. The question is whether the reduced information that results from projecting the complexity of epigenetic control into the reduced space of overall compaction, suffices for a biophysical description. Expanding this epigenetic potential *V* (*δ*Ψ) about a local minimum led to the results described in this paper. However, we should ideally think of this potential as itself evolving over some time scale and the choice of the initial point as reflecting a cell-specific initial condition, such as cellular levels of Lamin A (Swift et al., 2013).

The fluorescence anisotropy measurements of labelled histones in the embryonic stem cells state presented in Ref. (Talwar et al., 2013), coupled to confocal microscopy measurements of the nuclear dimensions, should provide a non-invasive way of determining the coupling of chromatin compaction to mechanical variables describing the nucleus and its shape. Examining other possibilities for simultaneously characterizing chromatin compaction in addition to nuclear size and shape in a non-invasive way would be especially valuable.

## Conclusions

In this paper, we presented a theory of auxetic behaviour in the nuclei of stem cells in the transitional state. We began by pointing out that the unusual mechanical properties of the stem cell nucleus, as well as its fluctuations, should provide a window into the packaging and dynamic character of the chromatin states contained within it. We argued that fluctuations in chromatin compaction should couple to fluctuations in the dimensions of the relatively soft nucleus that characterizes stem cells. We used these observations to argue that these coupled fluctuations, in chromatin packaging and nuclear shape, was most easily described in terms of a coupled, in general non-linear, dynamical system in three variables. We exploited the experimental observation that chromatin is least compact in the transitional state as compared to the pluripotent state and the differentiation primed state, to argue for a specific sign of the coupling term that connected size fluctuations to chromatin density fluctuations. We then showed how auxeticity resulted as a consequence, providing a simple and intuitive explanation for this puzzling observation. We then went on to suggest that we could map out the normal to auxetic transition using ideas from the model. We further suggested experiments that could implement and test these ideas.

We proposed that projecting the complex spatial-temporal distribution of chromatin compaction onto an overall compaction variable and interpreting the time-dependence of this variable in terms of motion within a simplified one-dimensional potential, should provide a particularly useful biophysical way of formalizing Waddington’s intuitive picture of an “epigenetic landscape” (Gilbert, 2000). This way of understanding landscape ideas in the differentiation of stem cells appears to be novel. Implementing the related analysis experimentally would seem to be feasible. Connecting microscopic, molecular-scale biochemical views of how stem cell transcriptional programs are modulated, with the averaged, larger-scale biophysical approach that we describe in this paper, should lead to an improved understanding of the determinants of stem cell mechanics and their coupling to chromatin states. This improved understanding should also help to illuminate the role of the mechanical environment in biasing lineage choice.

## Methods

### Derivation of model equations: R equation

Our equations are motivated in the following way, illustrated, for simplicity, in the isotropic case: Assume first that the nuclear is a sphere of radius *R*, prestressed by chromatin polymer pressure. Given compressibility, assume that the dominant modes of fluctuations are breathing modes, associated with an elastic energy 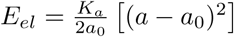, which penalises changes in area a from an unstressed or even pre-stressed state where the area is *a*_0_. This term also accounts for the contribution of the actin cytoskeleton, which enters as a modified area expansion modulus *K_a_*. Fluid flow in and out of the sphere, driven by a pressure imbalance, leads to volume changes and is resisted by a cost for deviations in the surface area from its preferred value. Describing stem-cell chromatin as a polymer solution at an effective (active) temperature *T*,* the free energy of the polymer solution in units of *k_B_T*^*\**^, is of the form 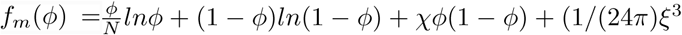 where ξ ∼ ϕ^−*v*/(3*v*–1)^ correlation term arising from monomer density fluctuations (Doi, 1996; Muthukumar, 2012). Activity enters as an effective temperature *T\**. More subtly, it modifies the effective Flory term *x*.

The polymer osmotic pressure follows from 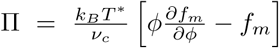, which yields 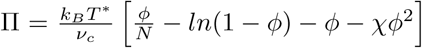, where *k*_*B*_ is the Boltzmann constant, *T*\* is the effective temperature, *v*_c_ is the monomer volume, *ϕ* is the volume fraction of the polymer and *N* is the degree of polymerization (Doi, 1996). Physically, *X* alters the relative balance of chromatin-chromatin and chromatin-solvent interactions, as manifest in the compaction state of chromatin. The effective Flory parameter *X* is then tuned by the fraction of bound nucleosomes, which controls Ψ: *x* = *x* (Ψ). We then have,

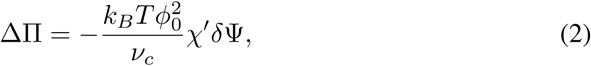

where 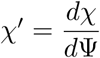. Penalising fluctuations of the nuclear envelope from its preferred area *a*_0_ yieldings a restoring net force of the form *F* = -16*πK_a_δR* and thus a pressure term

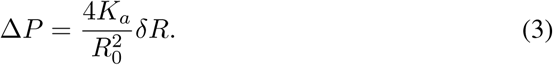

Darcy’s law provides an expression for the rate of change of volume 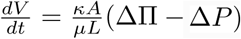, whereκis the permeability (m2), A is the area of the nucleus, *μ* the viscosity and L the length over which the pressure drops (Whitaker, 1986). This yields, where we use the notation 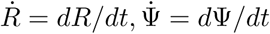

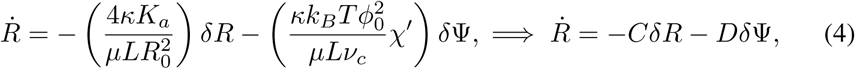

where 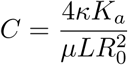 and 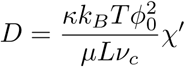. Note that *D* > 0 is required by the biophysical input that the binding of histones *must* lead to a contraction in DNA. The larger the polymer-solvent interaction, the smaller the Flory-Huggins *X* parameter implying that we can interpret histone binding and the consequent compaction of DNA as an effective decrease of the polymer-solvent interaction with histone binding. This then implies that the effective Flory-Huggins parameter should *increase* with Ψ, implying that *X*’ > 0. Here, ΔII *-*Δ *P* provides the driving force, in this case the difference between polymer and Laplace pressures relative to their unperturbed values. This holds in the absence of a force *f*. This result is easily generalized to the anisotropic case.

### Derivation of model equations: Ψ equation

We now discuss the dynamics of *δ*Ψ. First, ignoring the coupling to *R*_⊥_ and *R*_∥_, we model fluctuations in as relaxing in an over-damped manner to Ψ_0_. This dynamics explores the one-dimensional landscape defined through the effective potential *V* (*δ*Ψ), with Ψ_0_ at least a local minimum. Consider *N* nucleosome binding sites on a piece of DNA, in equilibrium with unbound nucleosomes at chemical potential *µ*, with the energy gain from nucleosome binding to DNA being Ψ. The probability of the nucleosome being bound is the Fermi function *p* = 1/(1 + e^(Ψ*-µ*)/k*B* *T*^). If the radius of the confining sphere is changed from *R*_0_ to *R* = *R*_0_ + *δR*, the DNA will stretch in place, altering Ψ. Assuming Ψ = Ψ(R), Ψ(R) = Ψ(R_0_ + *δR*) *π* Ψ(R_0_)+ Ψ^*’*^*δR* where 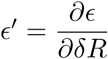. Expanding 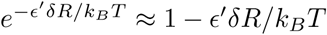, yields 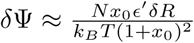 where 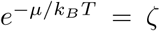 and 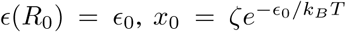. Thus, changes in *R* also drive changes inΨ, which evolves to its final value, given the change *δR*, through a kinetic coefficient which multiplies the term above. Adding to this the term in δΨ coming from the epigenetic potential, which can be assumed to be quadratic at lowest order in an expansion about the stable value 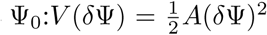, we have our final result: 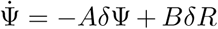 The sign of B depends on the sign of Ψ^*’*^, since all other quantities that enter its definition (*N*, *x*_0_ and *B*) are explicitly positive, reducing to the question of whether the nucleosome binding energy is *reduced* when the nucleus is expanded. In general, as is known from *in vitro* single molecule experiments, extending DNA expels bound nucleosomes, implying that their binding energy is reduced upon stretching; thus, the sign of *B* should be negative for the auxetic state given our interpretation above.

### Estimation of parameters

We now estimate 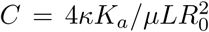 and 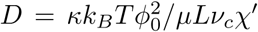. We take 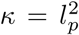, relating the permeability to the pore size *l*_p_. Assuming a nuclear pore complex size of *l*_p_ ≃ 5nm (Davis, 1995), this yields κ = 2.5 *×* 10^-17^m^2^. From plate theory, the area modulus *K_a_* and the Youngs modulus *E* are related through K_a_ ≈ *Et*, (Landau and Lifshitz, 1986) where *t* is the thickness of the plate. Thus, *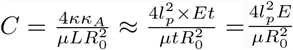*. With E ≈ 200 *Pa* (Caille et al., 2002; Guilak et al., 2000; Dahl et al., 2005), the radius of the nucleus *R*_0_ = *δ*Ψ *×* 10^-6^m and *µ* ≈ 2 - 3 centi-poise ≈ 2 *×* 10^-3^ Pa-sec, (Mastro et al., 1984) *C* ≈ 0.4sec^-1^. To calculate *D*, we assume that the length over which the pressure drops is the same as the membrane thickness (65 nm (Franke, 1970)). With *v*_c_ = (10nm)^3^ (Finch et al., 1977; Luger et al., 1997; Richmond and Davey, 2003), the polymer volume fraction ≈ 0.1 and *T* *≃* 300 K, we obtain D ≈ 8 *×* 10^-6^*X*^*’*^*m/sec*.

### Mapping to the experimental system

In steady state, 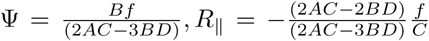 and 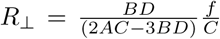. For finiteness, we require 2Cκ3D 蠀 0. From this, the Poisson’s ratio is 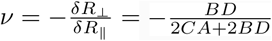. Choosing the experimental value of *v* = *-*0.25 and rearranging the above expression, we find that 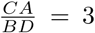 Making a reasonable choice for the ratio τ /τ_R_ *’* 0.01, yields τ_Ψ_ and the value of *C* obtained above yields τ_R_ = 2.5 sec and τ_Ψ_ = 2.5 *×* 10^-2^ sec, with A = 40sec^-1^. Fom 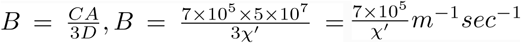,. Our final set of parameter values is then 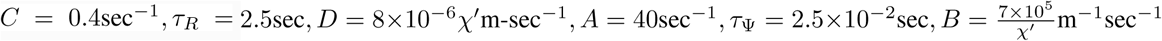. In dimensionless units, taking τ_Ψ_ = 2.5 *×* 10^-2^sec and measuring length in units of *R*_0_, yields: A = 1.0,C = 0.01,D = 0.04x^*’*^ and B = 0.09/x^*’*^. If we assume 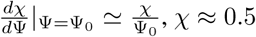 and Ψ_0_ = 1, this yields x^*’*^ = 0.5.

### Numerical simulations

Our numerical simulations implement Langevin dynamics in solving the stochastic Eqns. 1. We use both a simple Euler discretization as well as a fourth order Runge-Kutta method, checking that both gave essentially similar results.

### Declaration of Interest

The authors declare no competing interests.

## Acknowledgments

KT thanks Ankit Agrawal and Rishu Kumar Singh for useful discussions. Both authors thank M. Muthukumar for valuable input. GIM has benefited from conversations with J-F. Joanny and Maxime Dehan. GIM is also grateful for support from the Shastri Indo-Canadian Institute under the Shastri Mobility program.

## Author contributions

Research was designed by GIM. Both authors (KT and GIM) performed research and computations. KT plotted figures. Both authors contributed to writing the manuscript.

## Competing Interests

The authors declare no competing interests.

## Supplementary Information

### S1 Exact solution of the anisotropic case for a harmonic epigenetic potential in the absence of noise

Our governing equations represent a three-dimensional, coupled and, in general, nonlinear dynamical system. The choice of a harmonic epigenetic potential *δ*Ψ ^2^/2 and a constant force *f* yields a linear system of equations that can be written as,

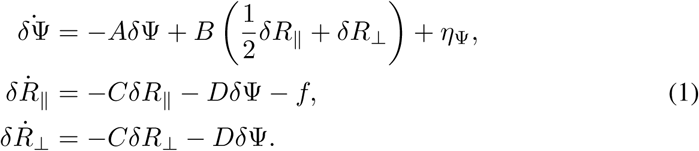

We define the Laplace transform and its inverse as,

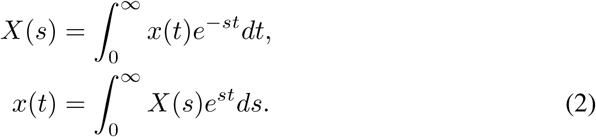

Using the definition in Eq 2, we take the Laplace transform of Eqs 1,

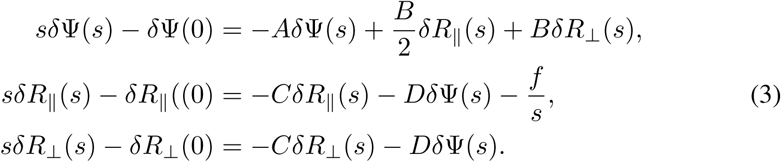

For the simplest initial condition, with *δ*Ψ (0) = *δR*_∥_(0) = *δR*_⊥_(0) = 0, taking the inverse Laplace transform yields the solution.

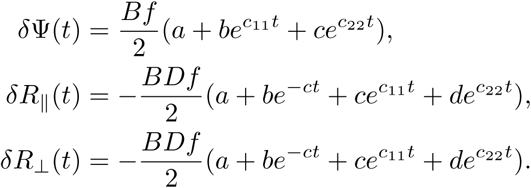

where the constants *a, b, c, c*_11_ and *c*_22_ are given by the following,

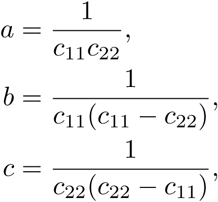

and

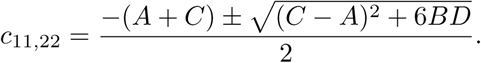

At long times, these solutions attain steady state values that vary linearly with the applied force f.

### S2 Correlation functions for the case of a harmonic epigenetic potential

For the system of equations which incorporates noise in the chromatin compaction variable, we can compute correlation functions analytically. We define Fourier and inverse Fourier transforms as:

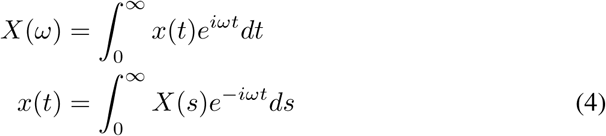

Taking the Fourier transform of the system and rearranging gives,

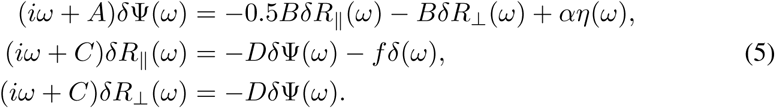

Solving this system of equations. Eqs. *δ*Ψ simultaneously, for *δ*Ψ (ω) yields the expression,

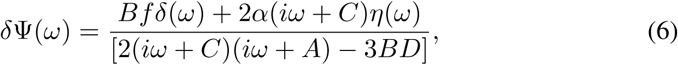

Using the expression for *δ*Ψ (ω) and the assumption that the noise is Gaussian and delta correlated yields

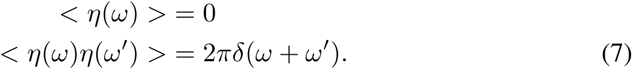

We thus obtain

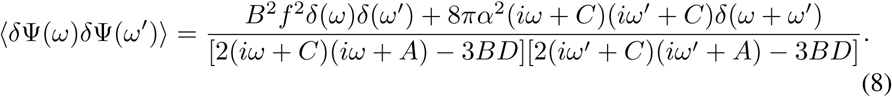

The inverse Fourier transform of the expression Eq. 8 yields the correlation function 〈*δΨ*(*t*)*δΨ*(*t*’)〉

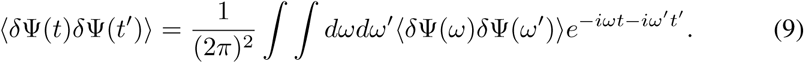

and

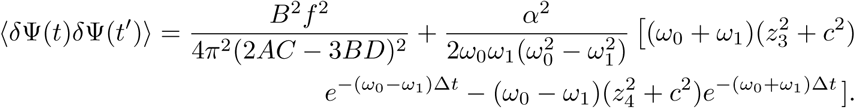

Now, we remove the constant part and keep the part coming from the fluctuation,

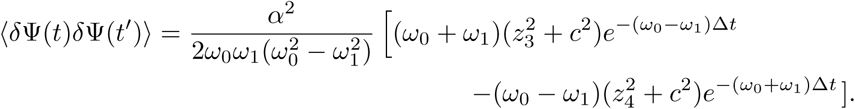

where Δ*t* = *t* - *t*^*’*^, *z*_1,2_ = *i*(*ω*_0_ ± *ω*_1_), *z*_3,4_ = *i*(−*ω*_0_ ± *ω*_1_), *ω*_0_ and *ω*_1_ is given by,

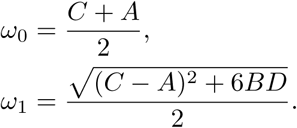

The other correlation functions are calculated in a similar way. The expressions are as follows,

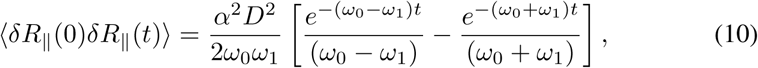

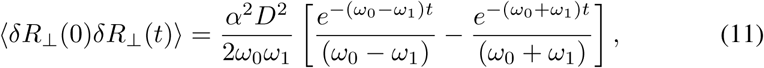

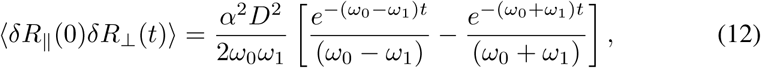

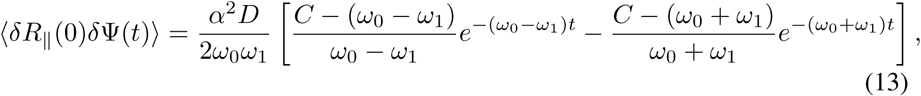

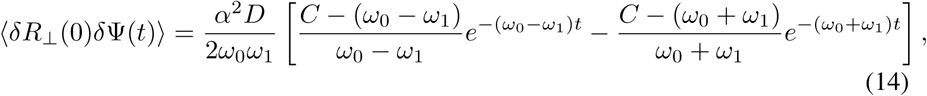

We display a number of autocorrelation and cross-correlation functions for the 3-dimensional auxetic system in S1.

**Figure S1:**
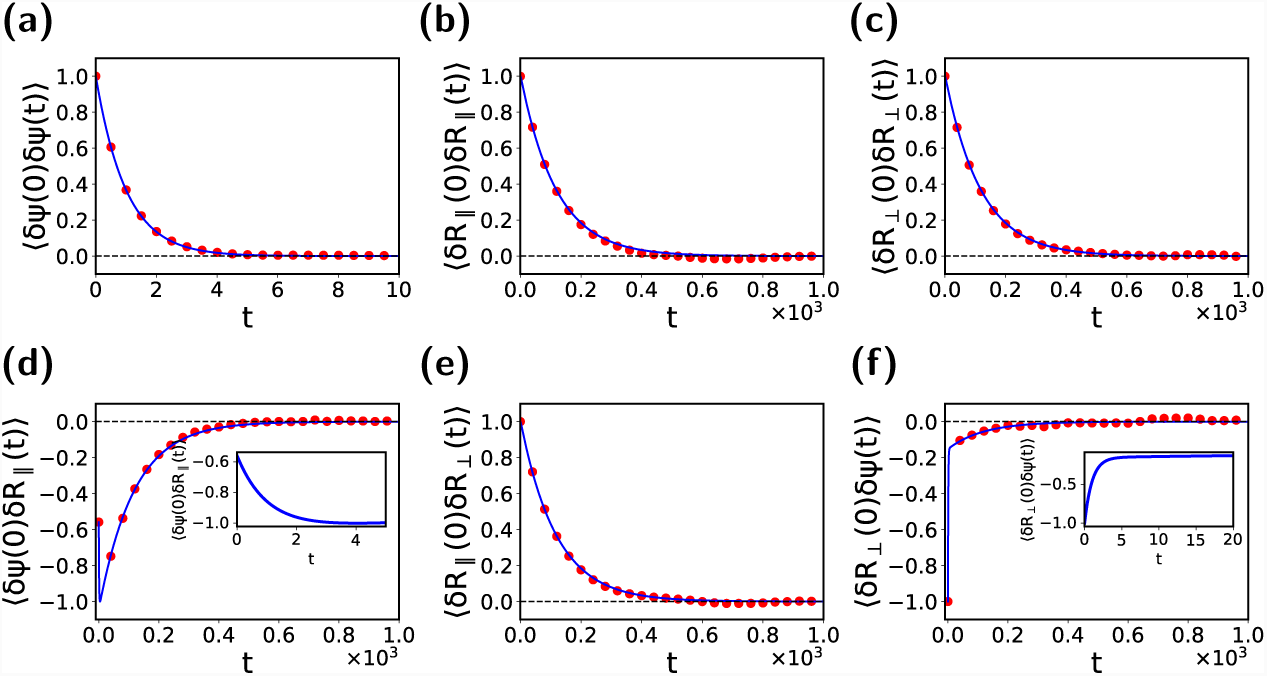
Computations of autocorrelations and cross-correlations, in the auxetic regime, with *C* = 0.01, *D* = 0.02 and *B* = *–-*0.17 **(a)** the autocorrelation function for *C, 〈c* (0)*c* (*t*)*〉*, **(b)** the autocorrelation function for *cR*_∥_, *〈cR*_∥_(0)*cR*_∥_(*t*)*〉*, **(c)** the autocorrelation function for *cR*_⊥_, *〈cR*_⊥_(0)*cR*_⊥_(*t*)*〉*, **(d)** the cross-correlation function for *C* and *cR*_∥_, *〈c* (0)*cR*_∥_(*t*)*〉*, **(e)** he cross-correlation function for *cR*_∥_ and *cR*_⊥_, *〈cR*_∥_(0)*cR*_⊥_(*t*)*〉* and **(f)** he cross-correlation function for *cR*_⊥_ and *C, cR*_⊥_(0)*c* (*t*). The insets show the behaviour close to the origin in two special cases where there is a competition between the two time-scales for relaxation. Points represent the numerical solution of the Langevin equations while lines represent the analytic formulae.

### S3 Long time behaviour of cross-correlation functions

The expression for the cross-correlation function *〈δR*_⊥_(0)δΨ (t)*〉* is,

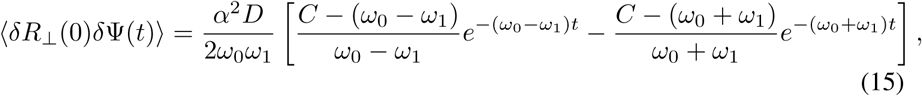

where ω_0_ = (*C* + *A*)/2 and 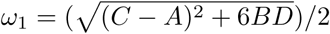.

From Eq. 15, it is evident that there are two time scales 1/(*ω*_0_ *ω*_1_) and 1/(*ω*_0_ + *ω*_1_). Since 1/(*ω*_0_ *ω*_1_) > 1/(*ω*_0_ + *ω*_1_), for the long term behaviour, we keep the *e*^-(*ω*_0_-*ω*_1_)*t*^ term and discard the *e*^-(*ω*_0_+*ω*1)*t*^ term. The resulting correlation function, in the long time limit, can be written as,

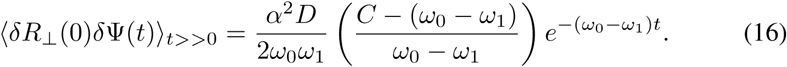

We normalize by,

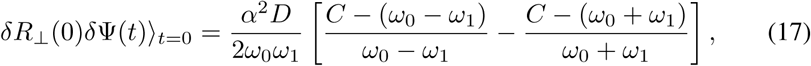

which yields,

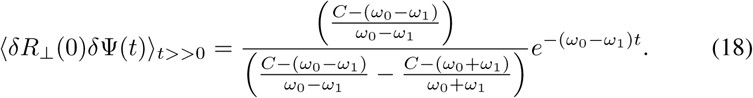

Eq. 18 for a choice of values of *B* is plotted in S2. The slope of the cross-correlation function *δR*_⊥_(0)δΨ (*t*) changes sign as the parameter *B* changes sign indicating the boundary of auxetic and non-auxetic regime. Similary the slope of the cross-correlation *〈δR*_∥_(0)δΨ (t)*〉* changes sign at the auxetic-nonauxetic boundary, while there is no such change in the slope of cross-correlation functions *δ*Ψ (0)*R*_∥_(t) and *δ*Ψ (0)*R*_⊥_(t) (Figure S3).

**Figure S2:**
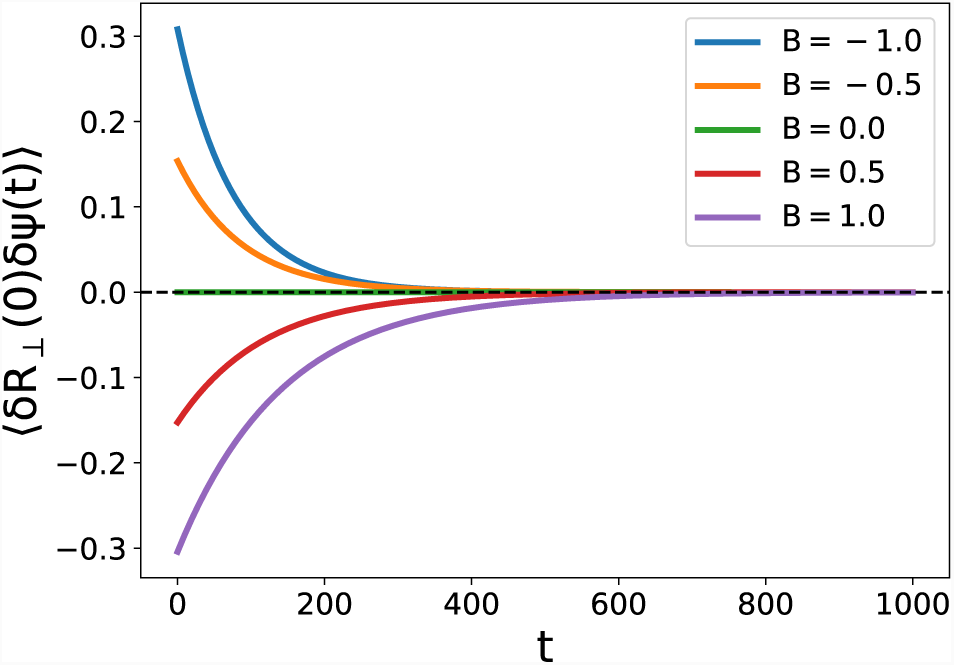
The figure shows the long time cross-correlation function *δR*_⊥_(0)*δΨ*(*t*) _*t>>*0_ with the various values of *B* from *B* = 1.0 to *B* = 1.0 including the *B* = 0 case. We see that the slope of the correlation finction changes as the sign of the parameter *B* changes from negative to positive.

The correlation functions *〈δR*_⊥_(0)δΨ (t)*〉* and *〈*δΨ (0)*δR*_⊥_(t)*〉* are same as *〈δR*_∥_(0)δΨ (t)*〉* and *〈*δΨ (0)*δR*_∥_(t)*〉*. Exchanging R_∥_ with R_⊥_ has no effect on correlation functions.

### S4 Periodic Force

The three-dimensional system with a harmonic potential and a periodic force can be written as,

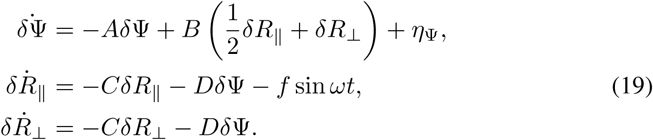

**Figure S3:**
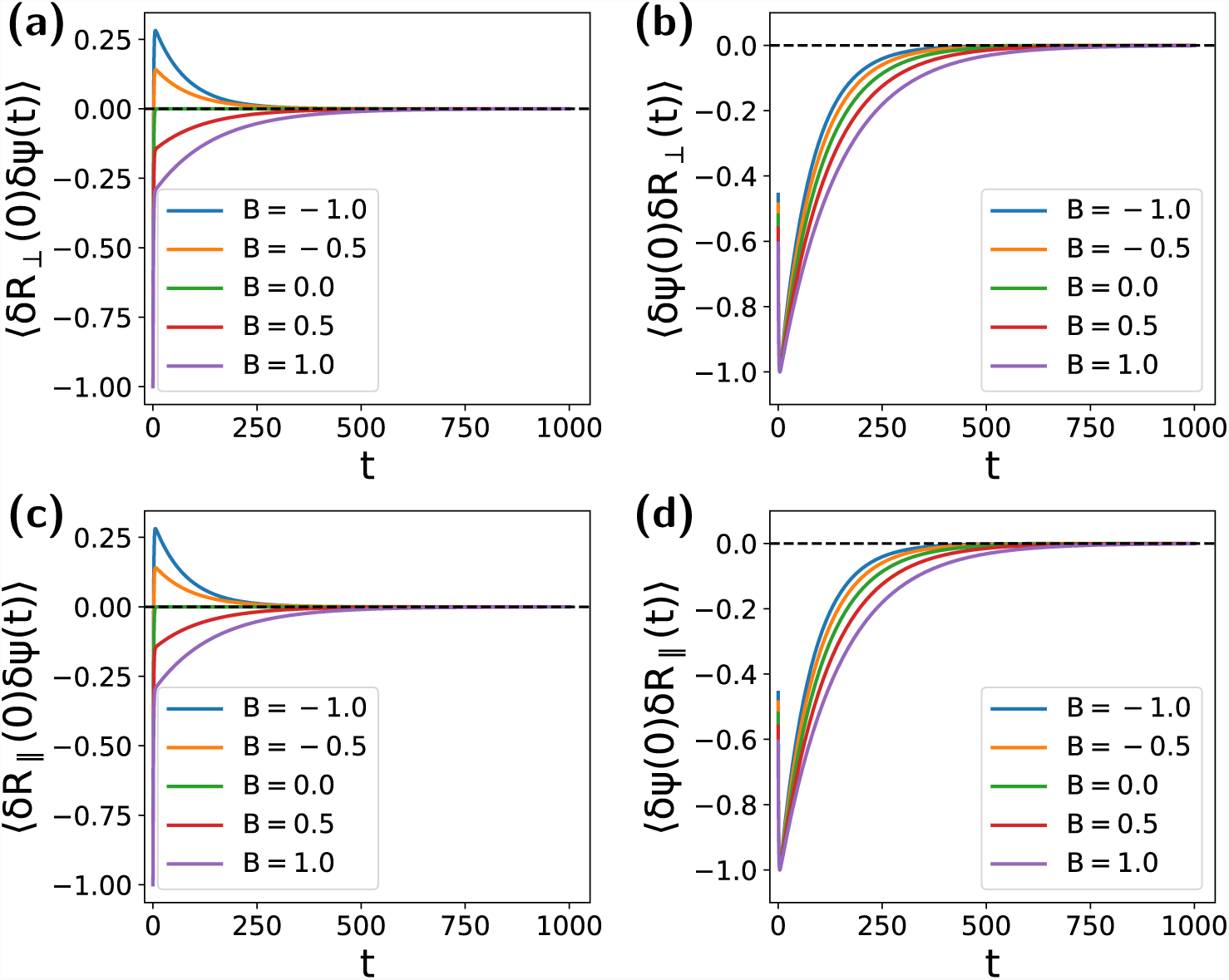
The correlation functions (a) the cross-correlation function for *δR*_⊥_ and *δΨ, 〈δR*_⊥_(0)*δΨ*(*t*)*〉* **(b)** the cross-correlation function for *C* and *δR*_⊥_, *〈δΨ*(0)*δR*_⊥_(*t*)*〉* **(c)** the cross-correlation function for *δR*_∥_ and *δΨ, 〈δR*_∥_(0)*δΨ*(*t*)*〉* **(d)** the cross-correlation function for *δΨ* and *δR*_∥_, *〈δΨ*(0)*δR*_∥_(*t*)*〉* with different values of parameter *B*. We see that the correlation functions *〈δR*_⊥_(0)*δΨ*(*t*)*〉* and *〈δR*_∥_(0)*δΨ*(*t*)*〉* change the slope with the parameter *B* while there is no such effect on *〈δΨ*(0)*δR*_⊥_(*t*)*〉* and *〈δΨ*(0)*δR*_∥_(*t*)*〉*.

Following the procedure of section S1,

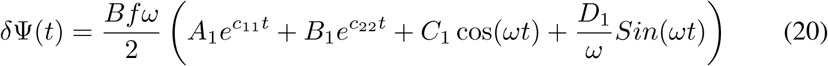

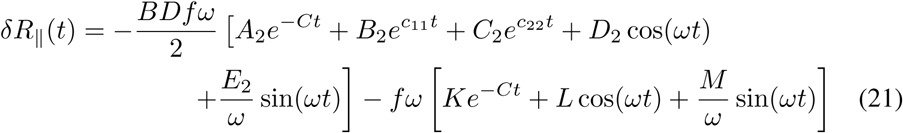

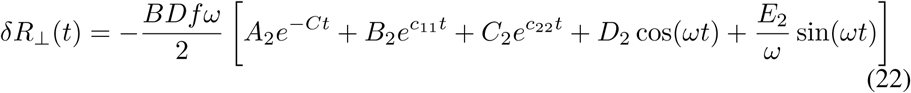

In order to get the steady state part of the solution, we discard the exponential terms which leads to the following expressions,

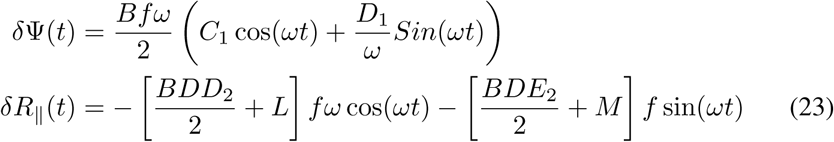

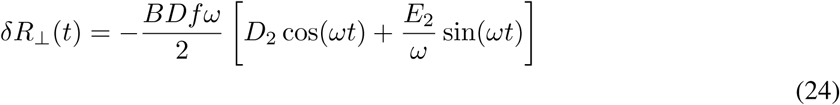

where the constants *A*_1_, *B*_1_, *C*_1_, *D*_1_, *A*_2_, *B*_2_, *C*_2_, *D*_2_, *E*_2_, *K*, *L* and *M* are given by the following,

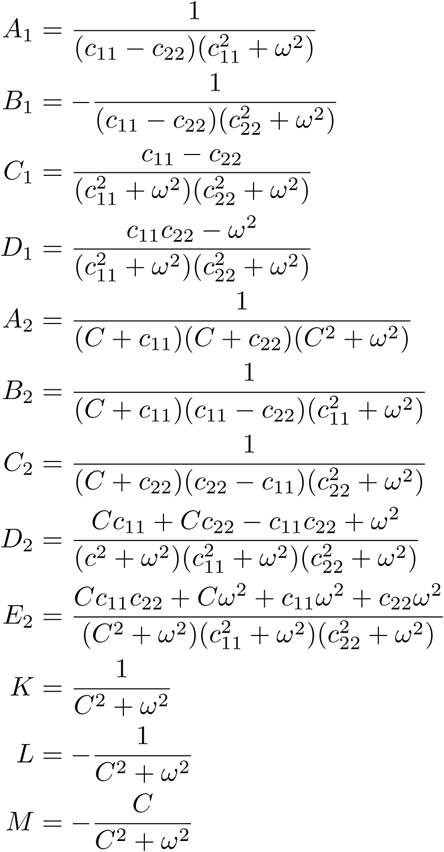

with 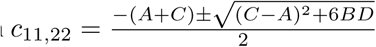

The variables *δ*Ψ (t), *δR*_∥_(t) and *δR*_⊥_(t) for auxetic and normal systems in steady state are plotted with time t (Figure S4).

**Figure S4:**
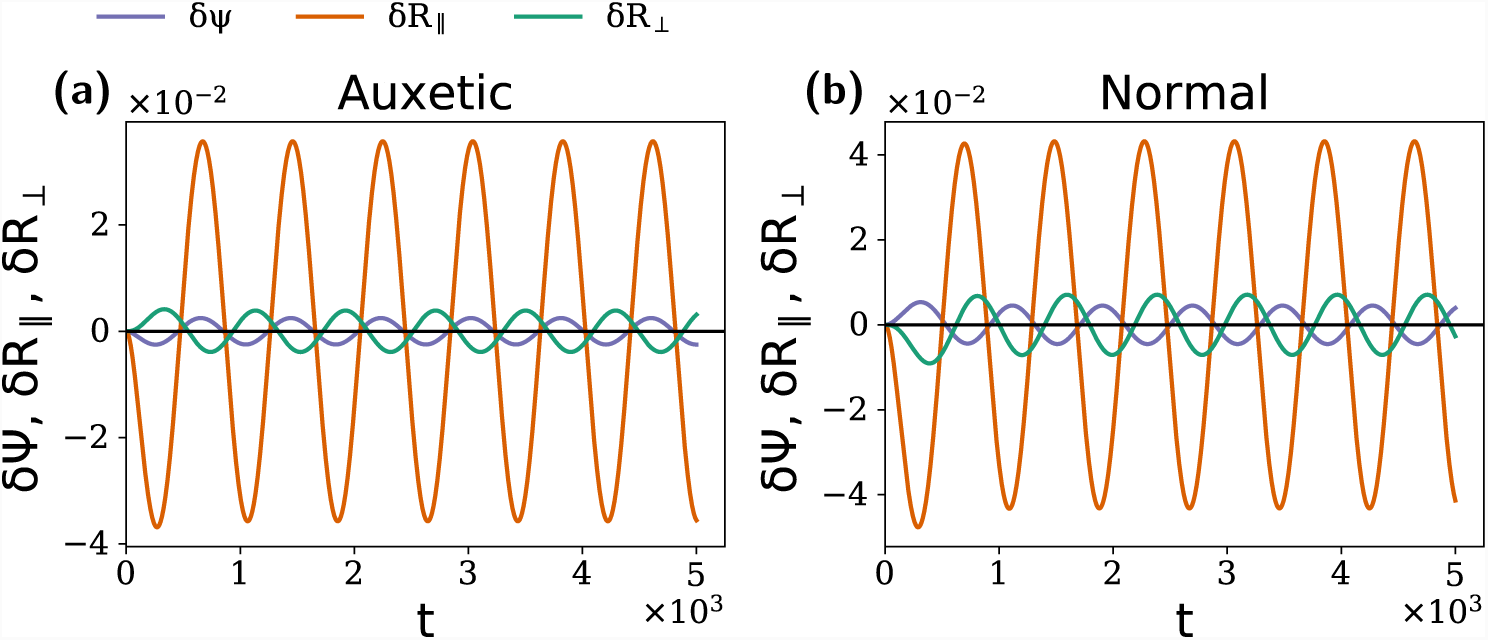
The behaviour of the systems under periodic force The figure **(a)** shows that how an auxetic system **(b)** a normal system behaves under a periodic force. It can be seen that in auxetic case, both radii *δR*_∥_ and *δR*_⊥_ simulanteously increase or decrease while in normal case, if one increases, the other decreases and vice-versa.

### S5 Reconstruction of the Potential Landscape

A simpler 2-d analog of the dynamical system (1) can be written as following,

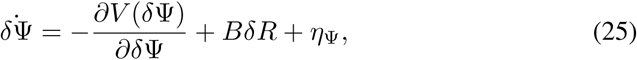

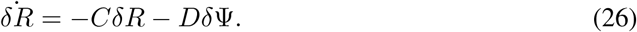

Assuming that the dynamics of *δR* is much slower than that of *δ*Ψ, we can consider *δR* as a constant in *δ*Ψ equation. This results in,

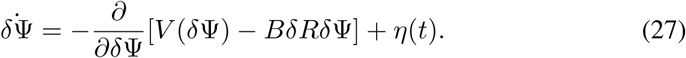

The corresponding Fokker-Planck equation can be written as,

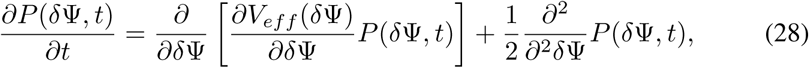

where *V_eff_* (*δ*Ψ) = *V* (*δ*Ψ) – *BδRδ*Ψ

For the steady state solution *∂P*/*∂t* = 0,

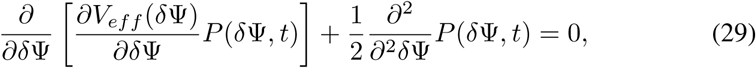

or,

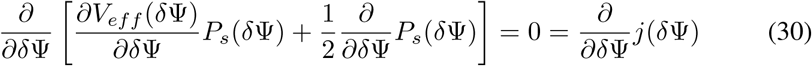

In steady state, the flux *j*(*δ*Ψ) vanishes, thus This means,

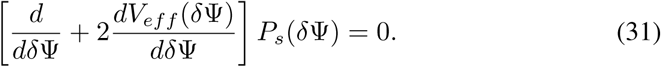

The above equation can be solved for the values of *δ*Ψ for a constant value of *δR*. Once we have those values, we can obtain the distribution *P* (*δ*Ψ). Taking the negative log of this result yields the effective potential *V_eff_*, as

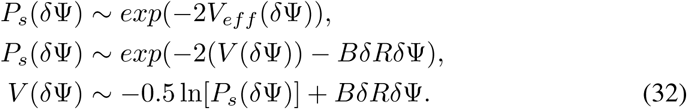

For the parameter value *B* = 0, this expression relates the probability distribution *P* (*δ*Ψ) to the potential landscape *V* (*δ*Ψ). This strategy is used in the reconstruction of the potential landscape in the Figure 6 of the main text.

